# A test for Hardy-Weinberg equilibrium on the X chromosome for sex-biased admixed populations

**DOI:** 10.1101/552794

**Authors:** Daniel Backenroth, Shai Carmi

**Affiliations:** Braun School of Public Health and community medicine, The Hebrew University of Jerusalem, Jerusalem, Israel

## Abstract

Genome-wide scans for deviations from Hardy-Weinberg equilibrium (HWE) are commonly applied to detect genotyping errors. In contrast to the autosomes, genotype frequencies on the X chromosome do not reach HWE within a single generation. Instead, if allele frequencies in males and females initially differ, they oscillate for a few generations towards equilibrium. Several populations world-wide have experienced recent sex-biased admixture, namely, their male and female founders differed in ancestry and thus in allele frequencies. Sex-biased admixture makes testing for HWE difficult on X, because deviations are *naturally* expected, even under random mating post-admixture and error-free genotyping. In this paper, we develop a likelihood ratio test and a *χ*^2^ test that detect deviations from HWE on X while allowing for natural deviations due to sex-biased admixture. We demonstrate by simulations that our tests are powerful for detecting deviations due to non-random mating, while at the same time they do not reject the null under historical sex-biased admixture and random mating thereafter. We also demonstrate that when applied to 1000 Genomes project populations (e.g., as a quality control step), our tests reject fewer SNPs (among those showing frequency differences between the sexes) than other tests.

## Introduction

Testing for deviations from Hardy-Weinberg equilibrium (HWE) is an important quality control step in genome-wide association studies ^1–4^. Extensive literature exists on HWE tests for the autosomes, from classic tests to recent work on Bayesian approaches, structured populations, sequenced or imputed genotypes, and software tools^5–18^. However, tests for HWE on the X chromosome have only been recently developed ^19–23^. The importance of associations of X-linked variants with complex traits, particularly as a mechanism of sexual dimorphism, has been recently recognized ^24–32^, and these developments underscore the importance of proper quality control on X, including testing for deviations from HWE.

A naive test for HWE on X would consider females only. However, such a test would implicitly assume an equal allele frequency between males and females. Indeed, a number of tests were recently proposed for *joint* testing of HWE in females as well as equality of allele frequencies between the sexes ^20–22^. However, these tests ignore the possibility that allele frequencies in males and females would differ *naturally* due to sex-biased admixture.

While autosomal allele frequencies reach HWE within a single generation, it is well known that for X, in case male and female allele frequencies initially differ, perfect equilibrium is never reached ^33,34^. The classical equations describing the evolution of allele frequencies on X, for an infinite population, are,

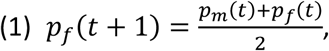

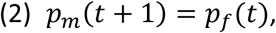

where *p*_*f*_(*t*) and *p*_*m*_(*t*) are the male and female allele frequencies, respectively, at generation *t*. Starting with unequal allele frequencies at generation *t* = 0, the male and female frequencies oscillate while gradually stabilizing. Specifically ^34^, if *p*_*f*_(0) = 1 and *p*_*m*_(0) = 1, then *p*_*f*_(*t*) = (2^*t*+1^ + (–1)^*t*^)/ (3 · 2^*n*^) and *p*_*m*_(*t*) = (2^*t*^ – (–1)^*t*^)/(3 · 2^*t*–1^). While equilibrium is approached exponentially quickly, if allele frequencies initially differ by a substantial amount, the frequency difference between the sexes can be non-negligible in the first few generations.

Recent sex-biased admixture has been known or identified for several populations, in particular in the Pacific and the Americas ^35–41^. Moreover, admixture in these populations has often been cross-continental, which may have led to large initial frequency differences between the sexes. Thus, even if a population has been randomly mating since admixture, and even if SNPs are accurately genotyped, we may expect *natural* frequency differences to exist for some X-linked SNPs, along with natural deviations from HWE in females. Thus, it would be wrong to discard X SNPs due to HWE violation, in case the violation can be explained as a natural result of sex-biased admixture.

In this work, we developed a likelihood ratio test and a *χ*^2^ test for HWE deviations on X, while permitting natural sex differences in frequency due to sex-biased admixture. This is achieved by taking into account the constraints imposed by Eqs. (1) and (2) on sex-specific frequency differences across generations. We show by simulations that our test has the expected size under the null, as well as power at least as high as existing tests for true deviations from the null (e.g., due to genotyping errors or inbreeding). Crucially, our test rejects HWE substantially less often compared to existing tests when HWE is violated due to historical sex-biased admixture in otherwise randomly mating populations. Finally, we show that in 1000 Genomes populations, our test rejects fewer SNPs among these for which frequency differences exist between the sexes.

## Methods

We denote the number of males and females in the sample as *n*_*m*_ and *n*_*f*_, respectively, and the two alleles as A and B. The numbers of male A and B carriers are denoted *m*_*A*_ and *m*_*B*_. The numbers of females with genotypes AA, AB, and BB are denoted *f*_*AA*_, *f*_*AB*_, and *f*_*BB*_. We denote by *p*_*m*_ and *p*_*f*_ the A allele frequencies in males and females, respectively.

We develop our likelihood ratio test based on the framework of You et al. ^21^. These authors have defined the *inbreeding coefficient ρ* to represent deviations from HWE. Using *ρ*, the expected genotype frequencies in females can be written as

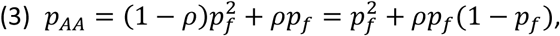

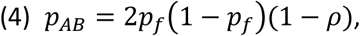

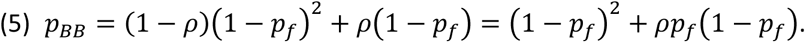

The null hypothesis of no deviations from HWE and no frequency difference between males and females is *p*_*m*_ = *p*_*f*_ = *p and ρ* = 0. We interpret here the parameter *ρ* more generally as a measure of the deviation from random mating in females, such that it can take any real value in [-1,1]. (This guarantees that all frequencies are in [0,1].) The alternative hypothesis is *p*_*m*_ ≠ *p*_*f*_ *or ρ* ≠ 0. Denote the parameters of the model as *θ* = (*p*_*m*_, *p*_*f*_, *ρ*). The likelihood of observing of the data (genotype counts) is multinomial,

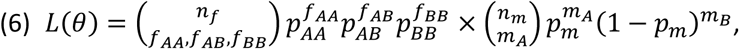

where *p*_*AA*_, *p*_*AB*_, and *p*_*BB*_ are given by Eqs. (3), (4), and (5), respectively. You et al. have proposed an expectation-maximization algorithm to obtain the maximum likelihood estimates (MLE) 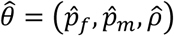.

Under the null hypothesis, *p*_*m*_ = *p*_*f*_ = *p* and *ρ* = 0, so *θ*_0_ = (*p, p*, 0), and the likelihood reduces to

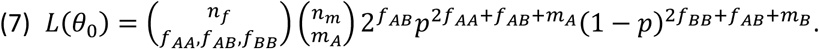

Here, the MLE is trivial, 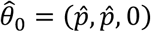, where 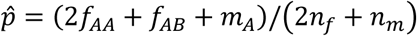. The likelihood ratio (LR) statistic is

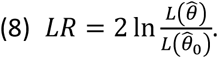

The LR statistic is then asymptotically distributed (under the null) as a *χ*^2^ distribution with two degrees of freedom, leading to a test we call the LRTP (likelihood ratio test for ***p***anmictic populations).

As explained above, the LRTP cannot accommodate “legitimate” frequency differences between the sexes due to sex-biased admixture. To address that, we reparametrize the model as follows. Instead of *θ* = (*p*_*f*_, *p*_*m*_, *ρ*), we write *θ* = (*p*_*f,g*_, *p*_*m,g*_, *ρ*), where *p*_*f,g*_ and *p*_*m,g*_ are the allele frequencies in females and males *in the previous* ***g****eneration*. With these parameters, the expected genotype frequencies in males in the current generation are

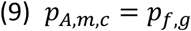

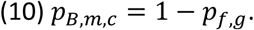

This is analogous to Eq. (1), which is true because males receive X chromosomes only from females in the previous generation. In females, assume for the moment that once *p*_*f,g*_ and *p*_*m,g*_ are given, females in the current generation are the products of random mating. The expected genotype frequencies in females in the current generation would be

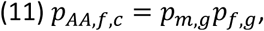

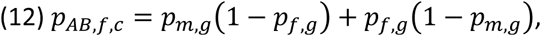

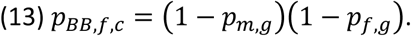

The above expressions reflect the fact that females receive one X chromosome from males and one from females. To incorporate deviations from random mating, we use again the parameter *ρ*. Analogously to the case of panmictic populations, we write the expected genotype frequencies in females in the current generation as

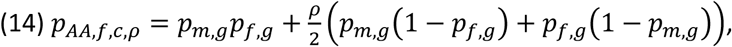

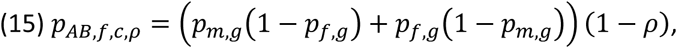

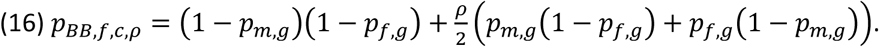

Note that the overall A allele frequency in females in the current generation is (for any *ρ*)

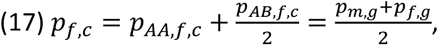

as expected based on Eq. (2). Also note that here, *ρ* cannot take any value, as *p*_*AA,f,c,ρ*_, *p*_*AB,f,c,ρ*_, and *p*_*AB,f,c,ρ*_ must all be within [0,1]. Our null hypothesis is that given the allele frequencies in the previous generation (*p*_*f,g*_ and *p*_*m,g*_), the genotypes of the current generation are determined by random mating, or *ρ* = 0. The alternative hypothesis is that there is a deviation from random mating, or *ρ* ≠ 0. The likelihood of the data under the most general *θ* is

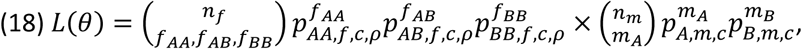

where *p*_*A,m,c*_, *p*_*B,m,c*_, *p*_*AA,f,c,ρ*_, *p*_*AB,f,c,ρ*_, and *p*_*BB,f,c,ρ*_ are defined by Eqs. (9), (10), (14), (15), and (16), respectively. The MLE 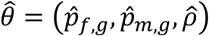 is obtained by taking the derivatives of (the logarithm of) *L*(*θ*) and equating to zero. This results in a set of three equations, which are too tedious to reproduce here, and can be solved numerically to yield the MLE 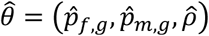. In practice, we directly maximized the log-likelihood based on a grid search. (We discarded any parameter set 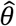 leading to allele frequencies in the current generation outside the range [0,1] in Eqs. (14), (15), and (16).)

In the case of random mating, *ρ* = 0, and thus the parameters are *θ*_0_ = (*p*_*f,g*_, *p*_*m,g*_, 0). The likelihood is

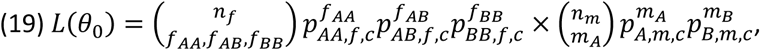

where *p*_*AA,f,c*_, *p*_*AB,f,c*_, and *p*_*BB,f,c*_ are defined by Eqs. (11), (12), and (13), respectively. Taking the derivatives of *L*(*θ*_0_) with respect to *p*_*f,g*_ and *p*_*m,g*_ and equating to zero results in the following pair of equations,

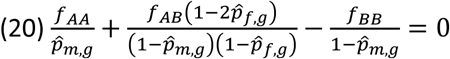

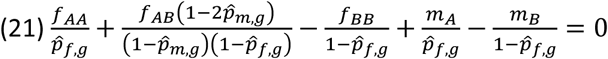

The solution of these equations yields the MLE under the null, 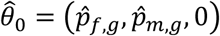. Here too, in practice we used a grid search to directly maximize the log-likelihood.

The likelihood ratio is then, as in Eq. (8),

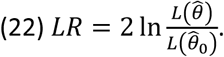

Under the null, LR is asymptotically distributed as *χ*^2^ with one degree of freedom, leading to a test we call LRTA (for ***a***dmixture).

For comparison, we also consider the LRTG test of ***G***raffelman and Weir ^22^. In their test, the likelihood of the data is as in Eq. (6), except that *p*_*AA*_ and *p*_*AB*_ are parameters to be estimated, and *p*_*BB*_ = 1 – *p*_*AA*_ – *p*_*AB*_ (i.e., *θ* = (*p*_*AA*_, *p*_*AB*_, *p*_*m*_)). The likelihood under the null is as in Eq. (7), and the likelihood ratio has a *χ*^2^ distribution with two degrees of freedom.

Finally, we also use our results to propose a new *χ*^2^ test ^22^. Suppose we have used Eqs. (20) and (21) to obtain the MLE 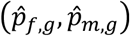. The expected values for the genotypes of males and females under the null (*ρ* = 0) are

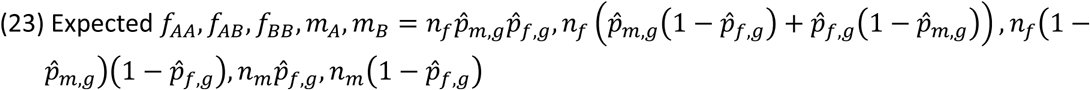

Then, given the observed values of *f*_*AA*_, *f*_*AB*_, *f*_*BB*_, *m*_*A*_, *m*_*B*_, a standard *χ*^2^ statistic can be calculated, which would be asymptotically distributed as *χ*^2^ with one degree of freedom. We call this test *χ*^2^-ML.

We also note that instead of the MLE 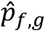 and 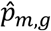, we could use a method of moments estimator, based on isolating *p*_*f*_(*t*) and *p*_*m*_(*t*) from Eqs. (1) and (2),

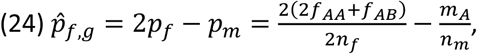

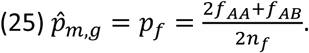

These estimates can then be substituted in Eq. (23), and a *χ*^2^ statistic can be calculated. We call this test *χ*^2^-MM. In practice, we found that the *χ*^2^-MM did not appropriately control the type I error rate (Table 1), and we did not report further experiments with that test.

**Table 1.** The proportion of rejections (Type I error rate) under random mating in a sex-biased admixed population. We compared the LRTP test (You et al. ^21^), the LRTG test (Graffelman and Weir ^22^), and the LRTA, *χ*^2^-MM, and *χ*^2^-ML tests developed in this paper. Our significance level was *α* = 0.05. The lowest proportion in each row is highlighted in bold.

| Number of generations since admixture | L RTP | L RTG | L RTA (this paper) | $\chi^2$-MM (this paper) | $\chi^2$-ML (this paper) |
| --- | --- | --- | --- | --- | --- |
| 1 | 1 | 1 | **0.05** | 0.22 | **0.05** |
| 2 | 0.97 | 0.97 | **0.04** | 0.09 | **0.04** |
| 3 | 0.47 | 0.48 | **0.06** | 0.07 | **0.06** |
| 4 | 0.17 | 0.18 | **0.04** | 0.05 | 0.05 |
| 5 | 0.09 | 0.09 | **0.06** | **0.06** | **0.06** |
| 6 | 0.10 | 0.10 | **0.06** | **0.06** | **0.06** |

## Results

We carried out several simulations to examine the behavior of our tests as compared to the LRTP and LRTG tests. We considered scenarios either under our tests’ null hypothesis, as well as under a number of alternative hypotheses.

Our first simulation was designed to examine the tests under their null hypothesis, namely sex-biased admixture with random mating thereafter. We started with a population of 400 males and 400 females, and a single locus with an initial allele frequency of 80% in females and 30% in males. Given the allele frequencies in one generation, we calculated the expected genotype frequencies in the subsequent generation based on Eqs. (9)–(13). Then, the genotypes of 400 males and 400 females were drawn based on multinomial distributions having these expected frequencies. We repeated the process up to six generations after admixture, and repeated the simulation 1000 times.

In Table 1, we report the proportion of rejections (type I error rate) when running five tests on the above genotype counts: the LRTP test of You et al. ^21^ and the LRTG test of Graffelman and Weir ^22^, both of which test for departures from either HWE in females or equality of allele frequencies between males and females; and the LRTA, *χ*^2^-ML, and *χ*^2^-MM tests we have developed here for sex-biased admixed populations (*Methods*). Our LRTA test and the *χ*^2^-ML test had an appropriate type I error rate (equal or close to the significance level *α* = 0.05), which is expected, because we simulated random mating post-admixture. In contrast, the LRTP and LRTG tests had much higher proportions of rejections, as expected due to the frequency differences between the sexes, which these tests are designed to detect. The type I error rate decreased to its value under the null (0.05) after about ≈6 generations post-admixture, when allele frequency differences between males and females became very small. The *χ*^2^-MM test did not control the type I error rate as well as the LRTA test and the *χ*^2^-ML tests, possibly because the parameters (allele frequencies in the preceding generation) are not accurately estimated. We thus do not further consider this test.

Our second simulation was designed to examine the power of the various tests under the alternative hypothesis of non-random mating. We considered one locus with an allele frequency of 80% in both males and females. We then calculated the expected genotype frequencies under one generation of mating, but this time with an inbreeding coefficient *ρ* equal to 0, 0.05, 0.1, 0.15, 0.2, 0.25, or 0.3, and simulated genotype frequencies in 400 females and 400 males based on the multinomial distribution with probabilities defined by Eqs. (9), (10), (14), (15), and (16). This simulation did not include sex-biased admixture, as the goal was to evaluate the power of our test under non-random mating, regardless of a history of admixture. We report the power of the various tests (at the 0.05 significance level and over 1000 repeats) in Table 2. The power of the *χ*^2^-ML test is always higher, followed closely by the LRTA test. The power of the LRTP and LRTG tests is slightly lower compared to our tests.

**Table 2.** The proportion of rejections (power) of the various tests under non-random mating of increasing strengths (without admixture). The highest proportion in each row is highlighted in bold.

| Inbreeding coefficient | L RTP | L RTG | L RTA (this paper) | $\chi^2$-ML (this paper) |
| --- | --- | --- | --- | --- |
| 0 | 0.04 | 0.04 | **0.05** | **0.05** |
| 0.05 | 0.12 | 0.12 | 0.16 | **0.19** |
| 0.1 | 0.37 | 0.37 | 0.49 | **0.52** |
| 0.15 | 0.69 | 0.70 | **0.82** | **0.82** |
| 0.2 | 0.92 | 0.92 | 0.95 | **0.97** |
| 0.25 | 0.99 | 0.99 | **1** | **1** |
| 0.3 | **1** | **1** | **1** | **1** |

Our third simulation was designed to validate that the LRTA and *χ*^2^-ML tests are powerful also under sex-biased admixture. We used the same approach as in our first simulation (Table 1), i.e., sex-biased admixture followed by random mating, except that after one generation, non-random mating was assumed with an inbreeding coefficient equal to 0.1, 0.2, or 0.3. We report the power of the LTRA and *χ*^2^-ML tests (at the 0.05 significance level and over 50 repeats) in Table 3. The power of the tests is unaffected by the historical admixture event (cf. Table 2).

**Table 3.** The power of the LRTA and *χ*^2^-ML tests in populations with sex-biased admixture and increasing strength of non-random mating.

| Inbreeding coefficient | LRTA<br>(this paper) | $\chi^2$-ML (this<br>paper) |
| --- | --- | --- |
| 0.1 | 0.48 | 0.49 |
| 0.2 | 0.98 | 0.98 |
| 0.3 | 1 | 1 |

Finally, we applied our methods to real data from the 1000 Genomes project ^42^ (1kG). We selected American populations in which sex-biased, cross-continental recent admixture was likely. While admixture in these populations has mostly ended 5-10 generations ago (e.g., ^43–46^), some SNPs may have not yet reached equilibrium, or were affected by more recent minor gene flow events. Our goal in this analysis was to determine whether our tests indeed reject less SNPs due to deviation from HWE. However, as the power of our tests was higher compared to the other methods (Table 2), the proportion of rejected SNPs may not be informative, since many rejected SNPs could be genuinely affected by genotyping errors. We considered instead a subset of SNPs where there was a significant evidence for allele frequency difference between the sexes, based on the test of Zheng et al.^19^, at P<0.05.

In Table 4, we report for each population the number of SNPs with a significant frequency difference between males and females, followed by the proportion of those SNPs rejected by each of the LRTP and LRTG tests as well as by our LRTA and *χ*^2^-ML tests. It can be seen that the proportion of rejected SNPs is lowest with our LRTA test. This result, along with the power simulations (Table 2), suggest that the LRTA test is likely to retain the maximal number of accurately genotyped SNPs for downstream analyses, while at the same time accurately detecting SNPs with true deviation from random mating. However, we note that in the absence of ground truth information on genotyping error status in 1kG, we cannot provide a formal proof of this claim.

**Table 4.** The proportion of rejected SNPs (at *α* = 0.05) under the LRTP, LRTG, and LRTA tests in 1kG populations. We restricted our comparison to SNPs with a statistically significant frequency difference between the sexes. The lowest proportion in each row is highlighted in bold.

| Population | #sex-biased SNPs | LTRP | LRTG | LRTA (this paper) | $\chi^2$-ML (this paper) |
| --- | --- | --- | --- | --- | --- |
| ACB (African Caribbeans in Barbados) | 34206 | 0.38 | 0.38 | **0.09** | 0.12 |
| ASW (Americans of African Ancestry in SW USA) | 33217 | 0.25 | 0.25 | **0.07** | 0.12 |
| CLM (Colombians from Medellin, Colombia) | 22815 | 0.40 | 0.4 | **0.11** | 0.16 |
| MXL (Mexican Ancestry from Los Angeles USA) | 12778 | 0.40 | 0.4 | **0.11** | 0.16 |
| PEL (Peruvians from Lima, Peru) | 14562 | 0.37 | 0.37 | **0.09** | 0.10 |
| PUR (Puerto Ricans from Puerto Rico) | 21455 | 0.41 | 0.41 | **0.11** | 0.12 |

## Discussion

In this paper, we proposed new tests for deviations from HWE on the X chromosome for sex-biased admixed populations. The X chromosome is unique in that allele frequencies do not reach equilibrium within one generation after perturbation, even when the population is otherwise randomly mating and all genotypes are observed without errors. Thus, the X chromosome requires a specialized test for HWE, even beyond accounting for the different ploidy between the sexes. Here, we proposed new likelihood ratio and *χ*^2^ tests to address this gap. We showed that our tests have the expected size (type I error rate) under sex-biased admixture and random mating thereafter, whereas other tests have high error rates, in particular when admixture was very recent. Additionally, our test has equal or higher power compared to the other tests considered. We also demonstrated that our tests reject fewer X chromosome SNPs in real 1000 Genomes populations. We thus recommend the application of our tests when performing quality control on the X chromosome. Our tests are available as an R package called HWadmiX at https://github.com/dbackenroth/HWadmix. Avenues for extending our approach can be the development of exact tests, Bayesian tests, or tests for multiple alleles.

## Acknowledgements

We thank Alon Keinan for discussions. S. C. thanks the German-Israeli Foundation for Scientific Research and Development (GIF) grant I-2489-407.6/2017 and the Israel Science Foundation (ISF) grant 407/17.

